# Demonstration of simultaneous biological sulphate reduction and partial sulphide oxidation in a hybrid linear flow channel reactor

**DOI:** 10.1101/569269

**Authors:** TS Marais, RJ Huddy, STL Harrison, RP van Hille

**Affiliations:** Centre for Bioprocess Engineering Research, Department of Chemical Engineering, University of Cape Town, Private Bag X1, Rondebosch, 7701, South Africa; The Moss Group, 13 Bell Crescent, Westlake Business Park, Westlake Drive, Tokai, 7945, South Africa

**Keywords:** Biological sulphate reduction, partial sulphide oxidation, semi-passive bioprocess, floating sulphur biofilm, linear flow channel bioreactor

## Abstract

Semi-passive remediation systems have the potential to treat low-volume, sulphate-rich, mining impacted waters in a cost-effective and sustainable way. This paper describes the “proof of concept” evaluation of a hybrid linear flow channel reactor, capable of sustaining efficient biological sulphate reduction and partial oxidation of the sulphide product to elemental sulphur. Key elements include the presence of a sulphate-reducing microbial community, immobilised onto carbon fibres and the rapid development of a floating biofilm at the air-liquid interface. The biofilm consists of heterotrophic species and autotrophic sulphide oxidisers. It impedes oxygen mass transfer into the bulk volume and creates a suitable pH-redox microenvironment for partial sulphide oxidation. Demonstration of the concept was successful, with near 20 complete reduction of the sulphate in the feed (1 g/l), effective management of the sulphide generated (95-100% removal) and recovery of a portion of the sulphur by harvesting the elemental-sulphur-rich biofilm. The biofilm re-formed within 24 hours of harvesting, with no decrease in volumetric sulphate reduction rate during this period. Colonisation of the carbon microfibers by sulphate reducing bacteria ensured biomass retention, suggesting the reactor could remain effective at high volumetric flow rates.

## 1. Introduction

Sulphide-rich mineral ore associated with gold and coal mining and related activities have left a legacy of acid rock drainage (ARD), which threatens the environment as well as ground and surface water in South Africa (McCarthy 2011). The impacted water is characterised by high levels of acidity, sulphates and potentially toxic metals with low concentrations of organic material (McCarthy 2011). The long-term nature of ARD generation, extending from decades to centuries, the impact on health and the ecology of receiving water bodies and the predicted increase in unmet water demand in SA drive the need for economically sustainable treatment options to address the ARD problem over extended periods of time.

The primary focus on remediation of ARD-contaminated water in South Africa has 40 been on high volume discharges emanating from abandoned underground mine basins, using established active technologies. Largely overlooked, the continuous, low volume ARD discharges from diffuse sources (waste rock dumps, discard heaps and open-pit mining) associated with coal mining, also have a significant impact on the environment due to the large number of sites and wide geographic distribution. Many are in rural or peri-urban areas, making traditional active treatment options unfavourable and passive or semi-passive approaches more favourable. Passive treatment options, such as wetlands, require less maintenance, fewer skilled operators and have lower operating costs. However, these systems (natural and constructed wetlands) are governed by slow, unpredictable kinetics and require extended hydraulic residence times (Skousen et al., 2017; Zagury et al. 2007; Sato et al., 2017), necessitating large land areas. Semi-passive ARD treatment systems present an attractive alternative for addressing these low-flow discharges, with lower capital and operational costs than active systems, improved control and greater predictability than conventional passive systems. There is an increasing interest in the development of semi-passive, sulphate reducing bioreactors as an alternative technology to traditional ARD treatment options at abandoned sites without electricity (Zhang and Wang, 2014).

Semi-passive systems, based on the action of sulphate reducing bacteria (SRB), can be applied to address sulphate reduction, metal precipitation and wastewater neutralisation simultaneously (Gopi Kiran et al., 2017). Under anaerobic conditions, 60 sulphate reducing bacteria (SRB) reduce sulphate, in the presence of a suitable electron donor, generating sulphide and alkalinity (Reaction 1) (Zhang and Wang, 2014).

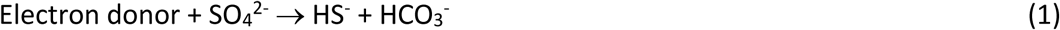

Despite extensive research demonstrating the technical feasibility and potential of biological sulphate reduction (BSR) for ARD treatment, relatively few commercial processes have been developed. The application of these technologies have been limited to niche applications, mainly due to the relatively slow kinetics of SRB, high cost of electron donor (e.g. ethanol, methanol and volatile fatty acids) and management of the sulphide product, which is significantly more toxic than sulphate (Rose 2013, van Hille et al., 2015). Each of these challenges must be addressed to develop a robust process that can be implemented.

Due to sulphide toxicity, the potential for re-oxidation, malodour, corrosivity and other hazards, an effective sulphide removal step is essential to ensure satisfactory treatment of sulphate-laden wastewaters (Jin et al. 2013, Harrison et al., 2014). Sulphur oxidising bacteria (SOB) produce elemental sulphur as an intermediate in the oxidation of hydrogen sulphide to sulphate under oxygen limiting conditions (Equation 2) (Kleinjan et al., 2005, Cai et al., 2017)).

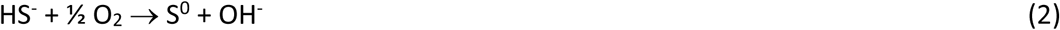

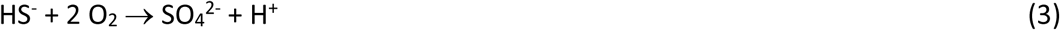

Sulphur produced by these microorganisms is stored in the form of sulphur globules, localised either inside or outside the cell (Cai et al., 2017). Biological sulphide oxidation has been applied in treating sulphide-rich waste streams (Cai et al., 2017), but commercial application has been limited to an active treatment approach, such as the Thiopaq® process (Janssen et al., 2000).

Biologically mediated partial oxidation of sulphide to elemental sulphur is restricted to a narrow pH and redox potential range (pH 6-8, −20 to −200 mV). In addition, the stoichiometric ratio of sulphide to oxygen needs to be maintained at 2:1 in order to facilitate partial oxidation to elemental sulphur and prevent complete oxidation to thiosulphate and sulphate (Reaction 3). This requires precise control of operational conditions, particularly the pH and supply of oxygen (Elkanzi, 2009).

Common drawbacks inherent with these active processes are the need for additional operational units and energy requirements, which increase capital and operational costs (Cai et al., 2017). A promising alternative is to control the partial oxidation of sulphide via a floating sulphur biofilm (FSB), a passive process that does not require energy input and provides an effective alternative for the removal of sulphide and recovery of elemental sulphur (Molwantwa et al.,2010; Rose, 2013). Floating sulphur biofilms were first observed on waste stabilisation ponds used to manage high sulphide tannery effluents. The biofilm impedes oxygen mass transfer, creating a discrete pH and redox microenvironment that facilitates partial oxidation of sulphide 100 from the bulk liquid, with deposition of the sulphur product in the organic biofilm (Molwantwa et al., 2010; van Hille and Mooruth, 2014).

The application of FSB for sulphide oxidation was first described in the Integrated Managed Passive IMPI process developed by Pulles, Howard and de Lange. Degrading packed bed reactors (DPBRs) were used for biological sulphate reduction, followed by linear flow channel reactor (LFCR) units for partial sulphide oxidation via FSB (Coetser et al., 2005). The process was evaluated at demonstration scale, but faced a number of challenges, particularly the sulphide oxidation component, which did not perform optimally (van Hille et al., 2011; van Hille and Mooruth, 2014). A detailed study by Mooruth (2013) led to further optimisation of design and operational parameters of the LFCR. The study demonstrated the feasibility of obtaining high partial oxidation rates in a sulphide-fed linear flow channel reactor (LFCR) through FSB formation.

Efficient BSR is dependent on maintaining a high biomass concentration and can be a challenge in flow through systems due to the relatively slow growth rate of anaerobic SRB. These limitations may be overcome by biomass retention. Support matrices for biomass attachment have been used to promote biomass retention in a number of reactor configurations to facilitate the decoupling of the hydraulic residence time and biomass retention time (Baskaran and Nemati, 2006; Sheoran et al., 2010, Hessler et al., 2018). A study by van Hille et al. (2015) demonstrated the potential application of carbon microfibers as support matrices for biological sulphate reduction within a 120 closed linear flow channel reactor (LFCR), providing a high surface area for biomass retention without significantly reducing effective reactor volume, as many bulky packing materials do. The study achieved a high sulphate reduction conversion of 85-95% at a feed sulphate concentration of 1 g/L. When operated at a dilution rate of 0.083 h_-1_, where substantial cell washout was observed in a continuously stirred tank reactor (CSTR) with a planktonic microbial community, the LFCR maintained a VSRR approximately 20% higher than the CSTR (van Hille et al., 2015). During the study, complete elimination of oxygen was not possible and there was evidence of partial sulphide oxidation and the establishment of a floating sulphur biofilm. This was similar to that observed in the dedicated sulphide oxidation reactor (Mooruth, 2013). This suggested that partial sulphide oxidation could be coupled with sulphate reduction within a single LFCR configuration.

This paper aims to build on the observations of Harrison et al. (2014) and van Hille et al. (2015) through the design and proof of concept evaluation of a hybrid reactor that integrates biological sulphate reduction and partial sulphide oxidation. It is hypothesised that the inclusion of carbon microfibres facilitates the attachment and retention of a sulphate reducing microbial community within the bulk volume of the reactor, while particular hydrodynamic properties (Mooruth, 2013) establish discrete anaerobic and microaerobic zones, the latter facilitated by the biofilm at the air-liquid interface. The biofilm creates a suitable microenvironment for partial sulphide 140 oxidation by sulphur oxidising bacteria. The paper assesses the performance of the hybrid channel reactor, in terms of sulphate conversion and volumetric reduction rate, sulphide removal and the potential for recovery of elemental sulphur.

## 2 Materials and methods

### 2.1 Microbial cultures

The mixed sulphate reducing bacteria (SRB) community used as a stock culture has been maintained at the University of Cape Town (UCT) over an extended period on modified Postgate B medium comprised of: 0.46 g/L KH_2_PO_4_; 1.0 g/L NH_4_Cl; 2 g/L MgSO_4_.7H_2_O; 0.3 NaSO_4_; 1 g/L yeast extract; 0.3 g/L sodium citrate and 1.6 ml of 60% sodium lactate (Oyekola et al., 2012). The synthetic feed was heat sterilised by autoclaving at 121°C, 103 kPa for 20 min. The sulphide oxidising bacterial (SOB) culture was developed at UCT using enrichments from SRB reactors (Mooruth, 2013).

### 2.2 Linear Flow Channel Reactor (LFCR) configuration and operation

A LFCR hybrid reactor (Fig. 1a.) was constructed from Perspex (11 mm thickness) with internal dimensions of 250 mm (l) x 10 mm (w) x 15 mm (h). The front facing side of the reactor was fitted with nine sampling ports, allowing the bulk reactor volume to be monitored across the length and at different heights. The reactor design (Fig. 1b and c) was based on the original 25 L LFCR described by Mooruth, (2013). The modified LFCR was fitted with a plastic strip (10 mm wide) holding carbon microfibers as a microbial support matrix and a heat exchanger (4 mm ID) for temperature control. A harvesting 160 screen, made of plastic mesh fixed to an aluminium frame, was designed to lie 5 mm below the liquid surface to facilitate biofilm harvesting. As illustrated in Fig. 1a, the feed was pumped into the LFCR continuously through the uppermost inlet-port on the left side of the reactor while the effluent flowed from the equivalent exit-port on the right side of the reactor. The reactor was set-up for the proof of concept evaluation at the 2.125 L scale.

**Figure 1:**
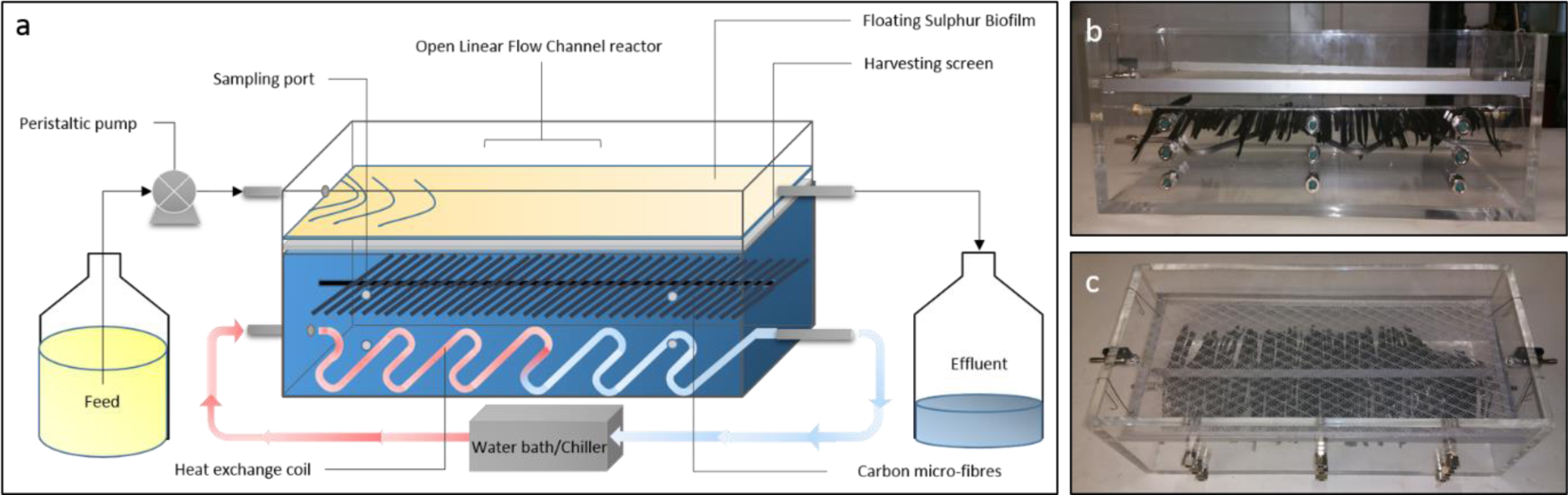
Images illustrating the LFCR configuration and design a) schematic diagram of the reactor set-up and operation b) side view and c) top view of the LFCR prior to inoculation showing the carbon microfibre strips, heat-exchanger and harvesting mesh.

The linear flow channel reactor (LFCR) was operated at a feed sulphate concentration of 1 g/L, supplemented with lactate as the sole carbon source to maintain a chemical oxygen demand (COD) to sulphate ratio of 0.7. While lactate is not a cost effective electron donor for large scale application it is extensively used as a preferred carbon source/electron donor for SRB research, promoting high biomass production and selecting for a diverse range of SRB species (Oyekola et al., 2010; Bertolino et al., 2012). The reactor was operated at 28°C with a hydraulic residence time of 4 days (dilution rate: 0.0104 h^−1^). The pH was not controlled.

### 2.4 Analytical methods

The pH was measured using a Cyberscan 2500 micro pH meter. Redox potential was recorded using a Metrohm pH lab 827 redox meter fitted with a Metrohm Redox platinum-ring electrode.

Dissolved sulphide was quantified using the colorimetric N,N-dimethyl-p- phenylenediamine method (APHA, 2012). Residual sulphate and thiosulphate 180 concentrations was determined by ion chromatography on a Thermo Scientific DIONEX ICS-1600 system equipped with an IonPac AG16 anion column, a 10 µl injection loop and a conductivity detector with suppression. A 22 mM NaOH solution was used as the mobile phase at a flow rate of 1 mL/min, as per manufacturers’ recommendations. Data were analysed using the Chromeleon®7 software package (version no. 7.2.1.5833).

The concentration of volatile fatty acids (VFAs) lactic, acetic and propionic acids in the feed and reactor samples was quantified using high pressure liquid chromatography (HPLC) on a Waters Breeze 2 system equipped with a Bio-Rad organic acid column (Aminex HPX-87H, 30 cm x 7.8 mm, 9 µm) and a UV (210 nm wavelength) detector. Acidified deionised water (0.01 M H_2_SO_4_) was used as the mobile phase at a flow rate of 0.6 mL/min, as per manufacturers’ recommendations (Biorad).

Elemental analysis of the floating sulphur biofilm was determined using an Elementar Vario EL Cube Elemental Analyser, for quantifying carbon, hydrogen, nitrogen and sulphur content of the sample (CAF, Stellenbosch University, South Africa).

Scanning electron microscopy (SEM) and energy-dispersive X-ray spectroscopy (EDS) was performed at the University of Cape Town Imaging Unit. Samples of FSB and colonised carbon fibres were fixed using 2 mL cold 2.5% glutaraldehyde in phosphate-buffered saline (PBS) solution (pH 7.2) for 24 hours at 4°C. After the primary fixation, the samples were washed twice with PBS solution, followed by an ethanol dehydration 200 series of steps using increasing concentrations (30, 50, 70, 80, 90, 95 and 100% (v/v)). Samples were mounted onto SEM stubs and critical point dried using hexamethyldisilazane (HMDS). The dried samples were coated with gold-palladium (60:40) and viewed using a FEI NOVA NANOSEM 230.

### 2.5 Hydrodynamics – Dye tracer study

The LFCRs were constructed from clear Perspex to allow for easy visualisation of hydrodynamic mixing patterns. A dye tracer experiment was conducted by filling the LFCR with a 2 mM sodium hydroxide solution to which ten drops of phenolphthalein was added to achieve a uniform pink colour. Hydrochloric acid (42 mM) was pumped into the reactor at predetermined flow rates. When the neutralisation reaction occurred (Equation 3), the liquid within the reactor turned colourless, demonstrating the fluid path. The experiment was performed at ambient temperature (22.4°C) across a range of hydraulic residence times (HRT) (4, 3, 2, 1, and 0.5 day). Each experiment continued till the entire reactor fluid volume turned colourless. This was monitored photographically, at intervals, throughout each experimental run (van Hille et al., 2011).

### 2.6 Proof of concept of simultaneous biological sulphate reduction and partial sulphide oxidation in a single reactor

To test the proof of concept, a 2 L LFCR was inoculated with a mixture of the SRB and SOB cultures and fed modified Postgate B medium containing 1 g/L SO_4_^2-^ using a speed 220 controlled peristaltic pump. The flow rate was equivalent to a 4 day HRT (dilution rate: 0.104 h^−1^). The temperature was controlled at 30°C. Samples (2 mL) were removed daily from the middle and lower sample ports in the first and third columns (FM, FB, BM, BB), as well as from the effluent port. The pH and sulphide concentration were measured immediately. The remainder of the sample was prepared for chromatographic analysis (VFAs and anions). Biofilm formation was observed visually. Once a thick, stable biofilm had been formed, it was collapsed and harvested periodically.

The FSB was collapsed by physically disrupting the biofilm and allowing the fragments to settle onto the harvesting screen. The sulphur product was recovered by removing the harvesting screen and collecting the accumulated biofilm. The biofilm was dried at 37°C for 48 hours, prior to weighing and elemental analysis.

### 2.7 Effect of biofilm disruption

To assess the effect of biofilm disruption, the reactor was operated for three residence times after which the floating sulphur biofilm (FSB) was broken up. Following disruption, the recovered FSB sample was collected and dried for elemental analysis. The reactor performance was monitored hourly for 24 hours after FSB disruption. The reactor was run for an additional three residence times before the biofilm was harvested.

## 3 Results and discussion

### 3.1 Hydrodynamics of the LFCR

A phenolphthalein tracer study was conducted to evaluate the fluid mixing profile in the LFCR (Fig. 2). The tracer study showed that the mixing in the LFCR was governed by the feed velocity (advective transport) at the the reactor inlet, which caused some short-lived, localised turbulent eddies. The absence of turbulent mixing and a slight density difference caused the acid feed to sink to the bottom of the channel. A dead zone occurred in the front corner of the reactor. The acid front moved along the floor of the reactor with a laminar parabolic profile (Fig. 2 b and c). After 30 minutes (Fig. 2b), the acid front reached the back wall of the reactor, resulting in the vertical displacement of the HCl layer. As time progressed, convective transport (a combination of advective and diffusive transport) became predominant (Fig. 2d and e), from the front and back of the reactor towards the middle, until the entire bulk volume turned colourless (macromixing time). This observed mixing profile in the LFCR was consistent across all HRTs tested.

**Figure 2:**
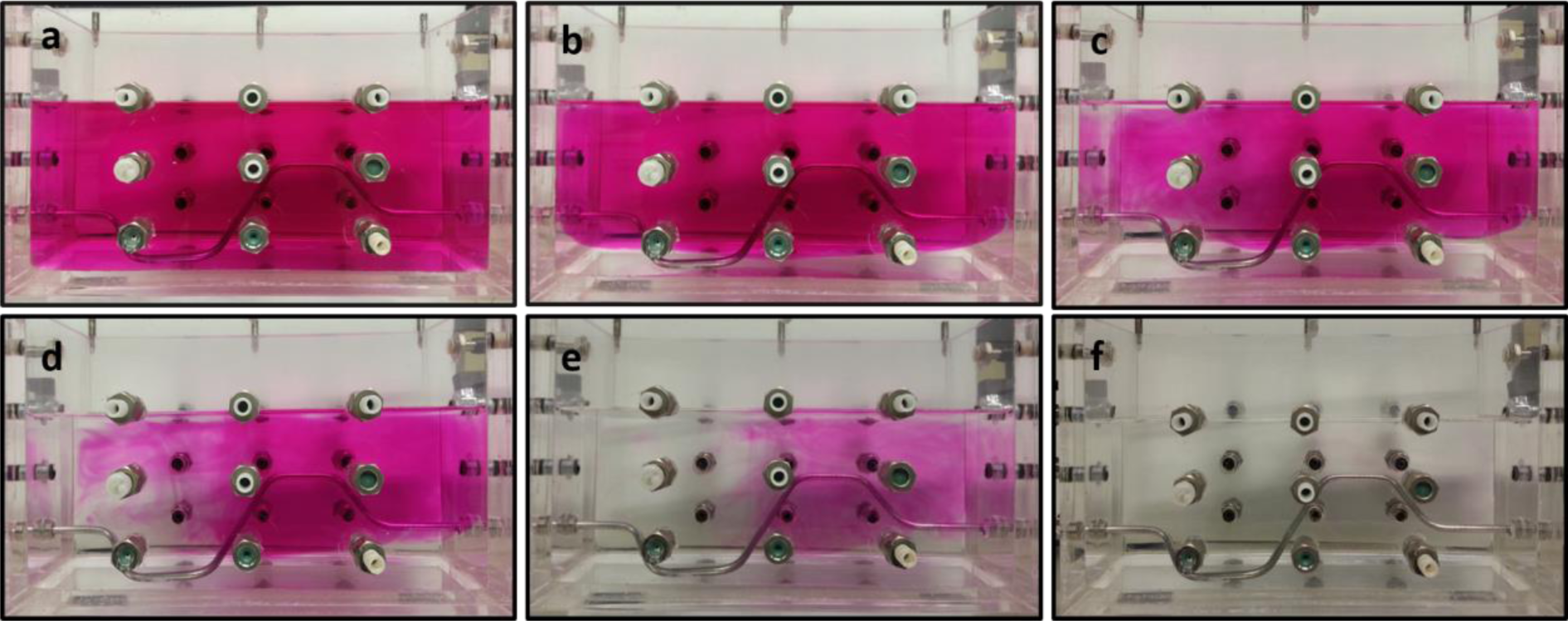
Dye tracer images showing the hydrodynamic pattern over time in a 2L LFCR operated at a 2 day HRT. a) t=0 b) t=30 min c) t= 48 min d) t= 1 h 54 min e) t= 2 h 15 min f) t= 2 h 25 min

The study concluded that the fluid dynamics in the LFCR is primarily governed by passive mixing through a combination of advective and diffusive transport. The relatively slow linear velocity and absence of turbulent mixing meant there was minimal disturbance at the surface of the reactor. Previous studies (Mooruth, 2013) calculated Reynolds numbers of less than 30, indicating very low levels of turbulence. This is critical to ensure suitable conditions to support the development of a floating 260 sulphur biofilm at the air-liquid interface. In addition, the current study revealed that the LFCR achieved macromixing times that were significantly shorter (2h 25 min) than the overall hydraulic residence times (2 day) tested. These findings are consistent with the conceptual fluid dynamic model previously described by Mooruth (2013). Though complete mixing times are significantly longer when compared to an active mixing system such as a CSTR. Results obtained from a saline tracer study, undertaken by measuring conductivity, revealed that the LFCR exhibited a residence time distribution (RTD) profile resembling that of a CSTR, due to the high diffusive mixing occurring within the bulk volume. This further supports the findings that the LFCR can be considered a relatively well-mixed system under the current conditions.

The fluid dynamics observed in the LFCR demonstrated the suitability of the reactor design for the desired application as it facilitates complete mixing within the bulk volume of the reactor with limited turbulence at the surface, promoting ideal conditions for sulphate reduction and the formation of the floating sulphur biofilm at the air-liquid interface.

### 3.2 Demonstration of the hybrid channel reactor

The LFCR was inoculated with an active SRB mixed microbial culture with an initial sulphide concentration of approximately 7 mmol/L (230 mg/L). The sulphide 280 concentration decreased rapidly over the initial 24 h as a result of unimpeded oxygen mass transfer across the liquid surface, resulting in sulphide oxidation. Within 24 hours a thin, but complete biofilm was observed covering the entire surface. Once the biofilm had formed the dissolved sulphide concentration in the bulk liquid began to increase steadily (Fig. 3a), from around 0.5 mmol/L (16 mg/L) to 4.6 mmol/L (152 mg/L) by day 34. This was an indication of SRB activity, which was further confirmed by the decrease in residual sulphate concentration from 10.86 mmol/L (1042 mg/L) in the feed to 3.89 mmol/L (373 mg/l) (Fig. 3a and c) by day 34, corresponding to a sulphate conversion of 64%. On day 34 a controlled biofilm collapse and harvest resulted in the rapid decrease in sulphide concentration and slight increase in residual sulphate concentration. As the floating sulphur biofilm regenerated at the air-liquid interface the sulphide concentration increased to 3.2 mmol/L by day 45 after which the biofilm was disrupted again. As the biofilm develops and matures it acts as a barrier, impeding oxygen mass transfer across the air-liquid interface. Colonisation of the carbon fibres resulted in retention of the SRB community and by day 60 the residual sulphate concentration (Fig. 3c) decreased to 1.0 mmol/L (<100 mg/L) reaching a VSRR and sulphate conversion of 0.11 mmol/L.h (10.56 mg/L.h) and 96% respectively.

**Figure 3:**
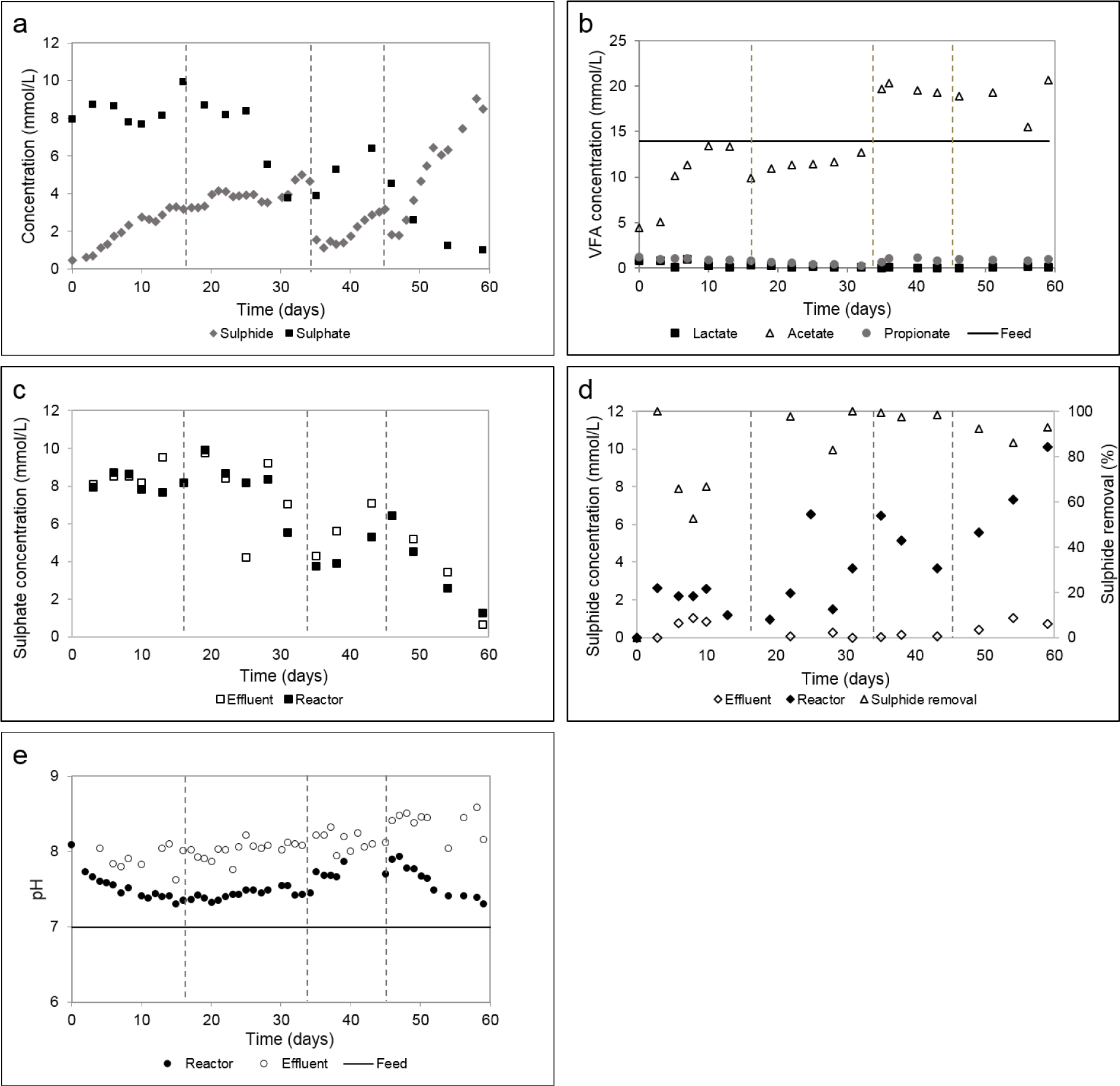
a) Residual sulphate and dissolved sulphide concentration profiles during the initial colonisation and stable performance phases of the hybrid LFCR b) measured volatile fatty acid concentrations c) reactor and effluent sulphate concentration d) 620 dissolved sulphide concentration and removal efficiency e) pH measured in the reactor and effluent over time during evaluation of the hybrid LFCR. Biofilm disruption events are indicated by vertical dotted lines.

The residual sulphate concentration (Fig. 3c) measured in the reactor and effluent was consistent throughout the study, indicating limited complete oxidation of sulphide. 300 Sulphide removal (Fig. 3d) was efficient, with low concentrations measured in the effluent. The average sulphide removal, between day 30 and 60, was over 90%. Thiosulphate remained below detection limits for the duration of the experiment. Together, these findings strongly suggest that partial sulphide oxidation to elemental sulphur was favoured throughout the study, with limited complete oxidation of sulphide to sulphate.

The LFCR maintained anoxic conditions within the bulk volume, with an average redox potential measured between −350 to −410 mV, an optimal range for sulphate reducing activity. The effluent samples were variable and exhibited increased ORP measurements. This was expected, since the effluent is exposed to the aerobic zone at the surface as it flows through the exit port of the reactor. The ability of the LFCR to maintain anoxic conditions within the bulk volume, despite the surface being open to atmosphere, was critical to achieve simultaneous sulphate reduction and partial sulphide oxidation.

The pH (Fig. 3e) increased initially in the bulk volume of the reactor (pH 7-7.5) with an additional increase observed in the effluent sample (pH 7.5-8). The initial pH increase was attributed to SRB activity as a result of alkalinity (bicarbonate) production while the additional pH increase observed in the effluent was attributed to partial sulphide oxidation, where hydroxyl ions are released as a by-product. While the experiments were conducted using feed at neutral pH, these results confirm the generation of 320 alkalinity and highlight the potential of the system to treat acidic wastewater streams.

The study assessed the anaerobic metabolism of lactate as a sole carbon source. The VFA concentration profiles were monitored over the period of the experiment. Lactate, a “high energy” substrate, can be metabolised via fermentation, oxidation or both by a wide range of microorganisms, shown in Reactions 5–7 (Bertolino et al., 2012, Oyekola et al., 2012):

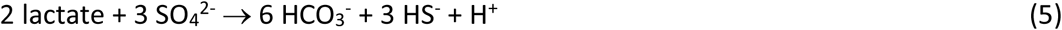

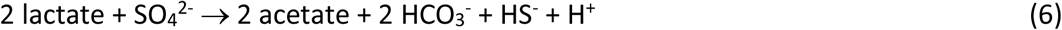

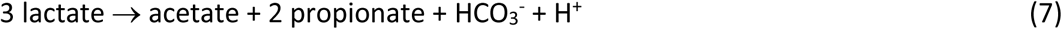

Throughout the experiment, the residual lactate concentration was below the detection limit. Acetate was accumulated as the main by-product, while propionate concentrations were consistently low. This indicates that incomplete lactate oxidation with concomitant sulphate reduction (Reaction 6) was the dominant metabolic pathway. The low propionate concentration suggests very little fermentation (Reaction 7) occurred throughout this study. These results were consistent with previous studies that investigated the VFA concentration profile in sulphate reducing reactors fed with lactate as a carbon source (Oyekola et al., 2012; Bertolino et al., 2012). Measured acetate concentrations exceeded the theoretical values based on reaction stoichiometry (amount of lactate supplied in the feed) from day 30 (Fig. 3b). This was attributed to the metabolism of yeast extract and citrate. Citrate metabolism is 340 described in literature, with acetate as a possible reaction product (Stams et al., 2009). Yeast extract is primarily added as a source of nitrogen, vitamins, and trace metals. However, it also contains carbon, in the form of carbohydrates (4-13%), which can be broken down to acetate (EURASYP, 2015). The accumulation of acetate means that it is unlikely that complete lactate oxidation (reaction 5) took place.

The study achieved a sulphate conversion of 96% with a corresponding VSRR of 0.11 mmol/L.h, when operated at a 4 day HRT. These results are consistent with data obtained using the same microbial community in conventional CSTRs under similar operating conditions (Oyekola et al., 2012). Between day 35 and 60 complete sulphide removal (95-100%) was achieved with the recovery of 30% of the added sulphur as elemental sulphur by harvesting the biofilm. A detailed study by Mooruth (2013) evaluated the chemical reactions that determine sulphur speciation in the LFCR and the dependence on pH and colloidal sulphur concentration. The study concluded that colloidal sulphur concentrations >2 mM in the pH range of 8.1 and 9.5 would result in polysulphide formation. Since the current system operated below pH 8 it is more likely that the fraction of elemental sulphur that was not recovered through the biofilm was suspended in solution. This was confirmed by HPLC analysis, which detected colloidal sulphur present in the effluent, as well as the accumulation of sulphur particles and biofilm fragments which settled in the effluent pipe and reservoir. This sulphur can be recovered by gravity sedimentation, while any formation of polysulphides downstream 360 can be rapidly converted to sulphur by decreasing the pH. This is a standard procedure applied in conventional sulphur recovery treatments (i.e. Thiopaq® process) (Cai et al., 2017).

Fig. 4 illustrates the development of the floating sulphur biofilm at the air/liquid interface over time. After 24 hours a thin layer of biofilm covered the surface, which continued to develop as the biomass and elemental sulphur content increased. The reactor configuration and operating conditions in the hybrid LFCR promoted the development and maintenance of two reactive zones. The conceptual model for the FSB was described by Mooruth (2013). Heterotrophic bacteria lay down an organic carbon matrix at the air-liquid interface, which supports the retention of autotrophic sulphur oxidisers. As the biofilm develops, it becomes a barrier to unimpeded oxygen mass transfer, creating a microaerobic zone within the biofilm, where pH and redox conditions favour partial oxidation of sulphide to sulphur. If sufficient sulphide is delivered from the bulk volume into the biofilm, complete consumption of oxygen is achieved and the bulk volume remains anoxic, favouring sulphate reduction. In this study, attachment of SRB to the carbon fibres supports biomass retention and increased sulphate reduction rates, ensuring that there is sufficient aqueous sulphide in the bulk volume to consume oxygen that penetrates in the time between biofilm harvesting and the establishment of a new biofilm. The harvesting screen is a critical component to the hybrid LFCR process serving as an effective mechanism for the 380 recovery of the floating sulphur biofilm with minimal disturbance to the process. It also permitted the biofilm to be disrupted intermittently, facilitating multiple cycles before a harvest is required (Fig. 4).

**Figure 4:**
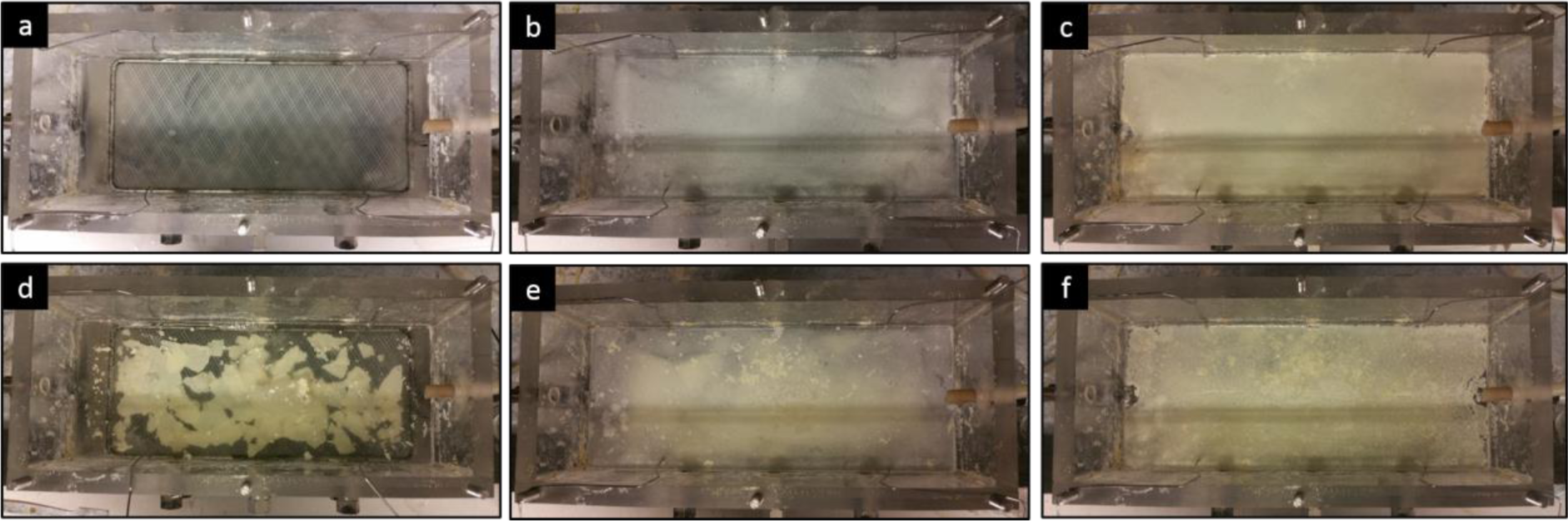
Progression of the floating sulphur biofilm formation at the air-liquid interface over time with a biofilm disruption event occurring on day 7. a) t= 0 b) t= 1 day c) t= 6 days d) t= 7 days (biofilm disruption) e) t= 8 days f) t= 11 days

### 3.3 Effect of biofilm disruption

Most BSR systems are operated in closed reactors under anaerobic conditions, due to the sensitivity of SRBs to oxygen. The disruption and harvesting of the biofilm removes the barrier to oxygen mass transfer into the bulk volume, until the biofilm re-forms. This could negatively affect the SRB activity, resulting in a decrease in performance or complete collapse of the system. Data from the initial demonstration (Fig. 3a) showed that FSB disruption had a significant effect on the sulphide concentration, with a notable decrease over 24 h. A study was conducted to evaluate the effect of biofilm disruption on reactor performance. This involved hourly monitoring of the reactor over the 24 h period after FSB disruption.

The biofilm was broken up on day 7. The removal of the biofilm resulted in a rapid decrease in dissolved sulphide concentration, from 4.5 mmol/L to 2.5 mmol/L within 12 hours, after which the concentration stabilised. A similar trend was observed in the effluent. Approximately 20 hours after disruption, the aqueous sulphide concentration began to increase again, corresponding with the reforming of the biofilm. After 24 hours a distinct thin layer of biofilm was observed (Fig. 4). The rapid decrease in dissolved sulphide concentration observed after biofilm disruption (Fig. 5a and b) can 400 be attributed to the increased oxygen mass transfer into the bulk liquid in the absence of the biofilm. The oxygen is rapidly consumed through the oxidation of sulphide. As the biofilm re-forms and matures, oxygen mass transfer into the bulk liquid is impeded and the rate of sulphide generation exceeds the sulphide oxidation rate, resulting in the observed increase in aqueous sulphide (Fig. 5a). Critically, the residual sulphate concentration remained stable during this period (Fig. 5b) indicating that sulphate reduction was not adversely affected during the 24 h period after biofilm disruption. It is clear that if there is sufficient residual aqueous sulphide to react with all the oxygen it is possible to maintain anoxic conditions in the bulk volume. This was confirmed by redox potential measurements.

**Figure 5:**
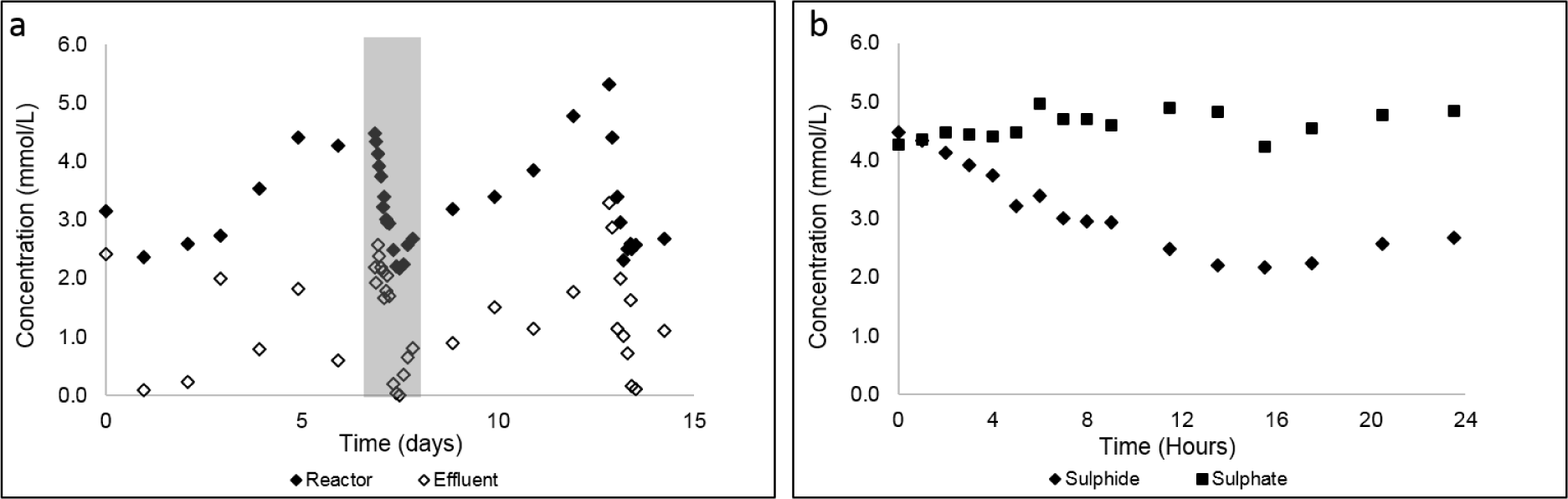
a) Effect of biofilm collapse on the performance of the hybrid LFCR showing a) dissolved sulphide concentrations in the reactor and effluent over 15 days and b) sulphate and sulphide concentrations in the bulk volume for the 24 h after FSB disruption (expanded view of the data highlighted in grey)

The decrease in sulphide concentration was predominantly attributed to sulphide oxidation rather than the evolution of H_2_S gas. This is consistent with the data from Mooruth (2013) who quantified H_2_S (g) liberation to account for mass balance discrepancies in a similar system. The study concluded that the liberation of H_2_S (g) from the LFCR surface was negligible. Most of the sulphide is present as HS_-_ at the operating pH and the lack of turbulent mixing reduces gas mass transfer across the surface. This is important from an aesthetic and safety perspective, due to the smell and toxicity of hydrogen sulphide gas.

Sulphide oxidation, with the generation of partially oxidised sulphur species such as colloidal sulphur, thiosulphate, polysulphides, or complete oxidation to sulphate may 420 occur abiotically, or catalysed by microbes. For abiotic oxidation, the thermodynamics associated with the initial transfer of an electron for sulphide and oxygen reveal that the reaction is unfavourable, as an unstable superoxide and bisulphide radical ion would need to be produced (Luther et al., 2011). Alternatively, a two-electron transfer is favourable, with the formation of a stable S_0_ and peroxide. However, the partially filled orbitals in oxygen that accept electrons prevent rapid kinetics. Due to these constraints the abiotic oxidation of sulphide is relatively slow. Alternatively, biologically mediated sulphide oxidation by photolithotrophic and chemolithotrophic microbes rely on enzymes that have evolved to overcome these kinetic constraints, allowing rapid sulphide oxidation. A study by Luther et al. (2011) demonstrated that biologically mediated sulphide oxidation rates are three or more orders of magnitude higher than abiotic rates. Furthermore, Mooruth (2013) investigated the extent of abiotic and biotic sulphide oxidation in a LFCR similar to that used in the current study. The study revealed that abiotic sulphide oxidation contributed little to the overall oxidation rate measured. This was based on a combination of kinetic constraints, poor oxygen diffusion, mixing (hydrodynamics) and convective mass transport within the LFCR. The biologically mediated oxidation of sulphide in the bulk volume following biofilm harvesting is critical. If the process relied on slower, abiotic oxidation, it is likely that the oxygen concentration in the bulk volume would increase to the point where sulphate reduction was inhibited.

As the biofilm develops, oxygen mass transfer into the bulk liquid is impeded and sulphide oxidation occurs exclusively within the biofilm. As the biofilm continues to mature and thicken, oxygen penetration through the biofilm slows to the point where it becomes limiting, resulting in significantly reduced partial sulphide oxidation and an increase in dissolved sulphide concentration in the effluent, which is undesirable. Therefore, there is a need to optimise the FSB harvesting frequency to ensure maximum sulphur recovery and consistent sulphide removal.

#### Scanning electron microscopy and energy-dispersive X-ray spectroscopy

The anaerobic and aerobic zones within the reactor promoted the development of two separate microbial communities. Examples of the different cell morphologies, visualised by SEM, are shown in Fig. 6. These images confirmed biomass attachment to the carbon microfibres.

**Figure 6:**
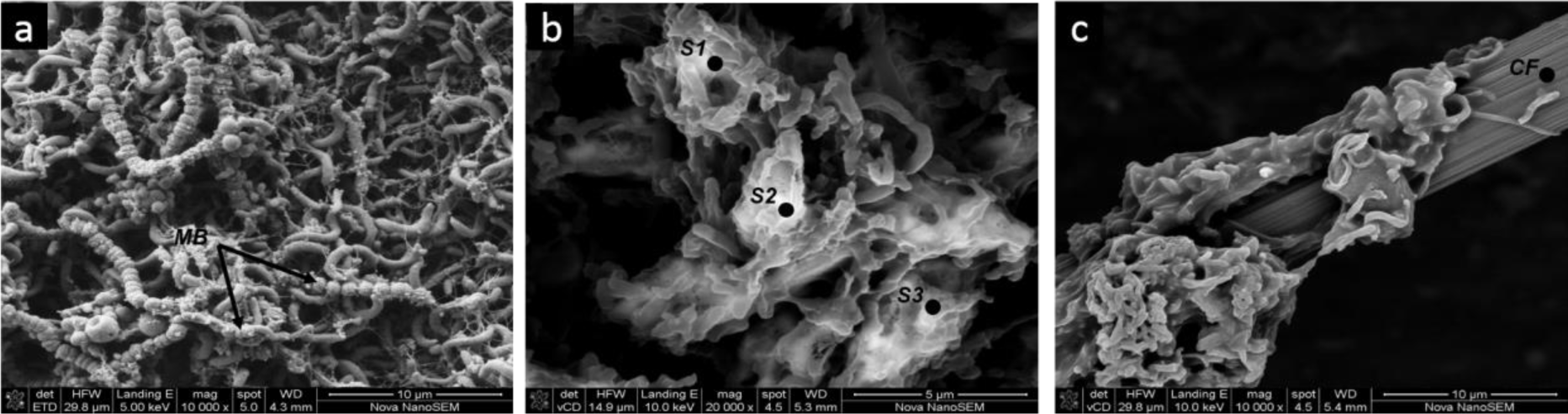
SEM images of the a) floating sulphur biofilm microbial community with membrane bound sulphur (MB) b) SEM-EDS imaging and elemental composition of the floating sulphur biofilm; point analysis of S1, S2 and S3 detected 100% sulphur (all elements were normalised) and c) attached biofilm on carbon fibre (CF)

Previous studies have reported on the enhanced VSRR that can be achieved through increased biomass retention within a LFCR fitted with carbon microfibres as the support matrix (van Hille et al., 2015). Additionally, SEM-EDS analysis (Fig. 6b) of the FSB confirmed partial sulphide oxidation, with the presence of highly concentrated (100%) elemental sulphur deposits detected across points selected shown on the image. Sulphur deposits were also observed on the outer membranes of the sulphur oxidising bacteria (Fig. 6a). This mechanism of sulphur excretion in SOBs has been previously documented (Cai et al., 2017).

### 3.4 Conceptual model for the hybrid LFCR

The data presented in this paper show that efficient biological sulphate reduction and partial oxidation of sulphide to sulphur can be achieved in a single reactor that is open to the atmosphere. The reactor was able to maintain sulphate reduction rates equivalent to those achieved in active, stirred tank reactors using a simple reactor geometry and no energy input. A conceptual model for how this is achieved is presented below and summarised in Fig. 7.

**Figure 7:**
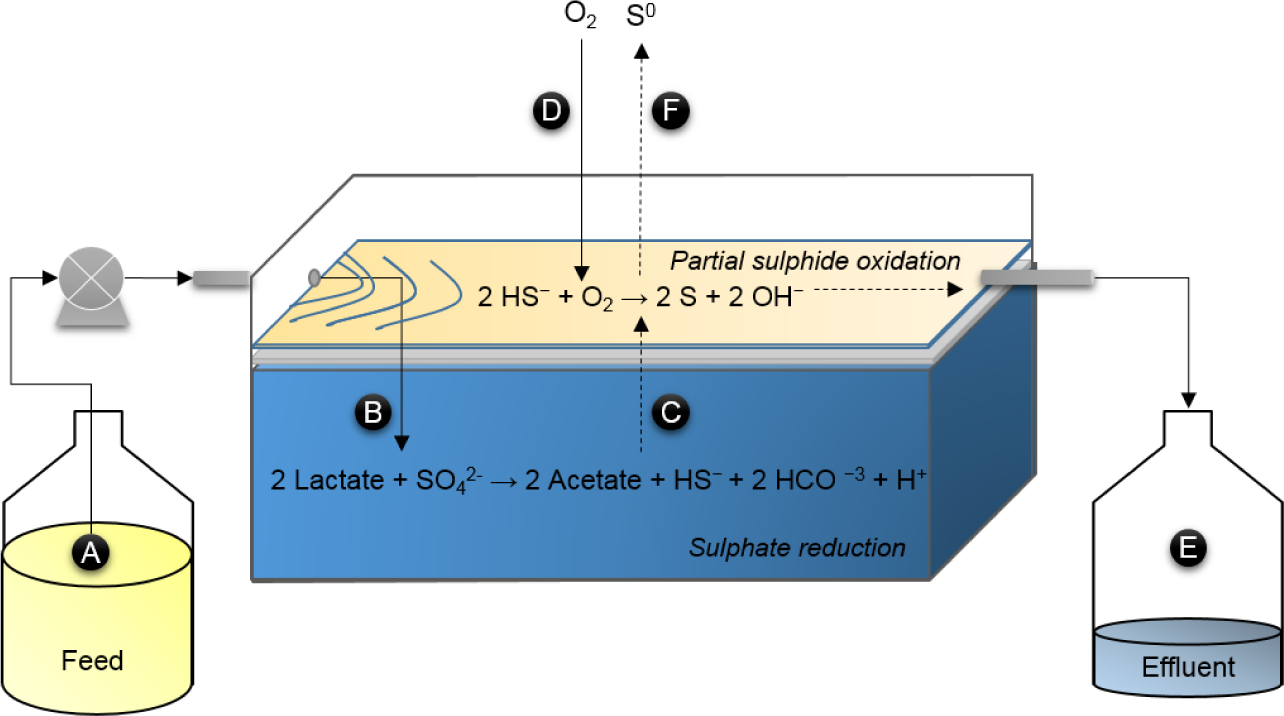
Conceptual model for the hybrid LFCR

Feed solution (A), containing sulphate and sufficient organic carbon to sustain sulphate reduction and the heterotrophic component of the biofilm is pumped into the reactor. Biological sulphate reduction occurs within the anaerobic bulk volume of the reactor (B). Microbial attachment and subsequent colonisation of the carbon fibres facilitates biomass retention and an increased sulphate reduction rate. The absence of turbulent mixing ensures limited loss of gaseous hydrogen sulphide, ensuring good odour control, while the fluid flow pattern ensures delivery of sulphide to the biofilm. The floating biofilm is formed, initially by heterotrophic species which produce an extracellular carbon matrix at the air-liquid interface. Autotrophic sulphur oxidisers colonise the biofilm. The biofilm results in an oxygen concentration gradient, creating a zone where the pH and redox environment favours microbially catalysed partial oxidation of sulphide to elemental sulphur. As the biofilm thickness and sulphur deposition increase, oxygen mass transfer is impeded to the point where the sulphide 480 oxidation rate becomes slower than the sulphide generation rate and the sulphide concentration in the effluent increases. At this point, biofilm disruption or harvesting is necessary. Disrupting the biofilm removes the barrier to oxygen mass transfer so oxygen from the atmosphere (D) diffuses into the bulk volume, where it is used by planktonic SOB to oxidise sulphide within the bulk volume. This leads to a rapid decrease in sulphide concentration, but critically ensures that the bulk remains anoxic and sulphate reduction is not inhibited. The biofilm begins to re-form almost immediately and within 24 hours oxygen mass transfer is reduced to the point where the aqueous sulphide concentration increases again. Treated effluent flows from the reactor (E) out of a port at the liquid surface and is characterised by low residual sulphate and sulphide. The biofilm is recovered by removing the harvesting screen (F) that lies just below the air-liquid interface.

## 4. Conclusion

This paper shows the successful integration of biological sulphate reduction and partial sulphide oxidation within a single LFCR unit, achieving near complete sulphate conversion (97%) to sulphide with associated partial sulphide oxidation under the conditions tested, leading to elemental sulphur recovery. Laboratory-scale demonstration of the hybrid LFCR is a first step toward a sustainable, low cost option for treating sulphate-rich, mining impacted water.

## Acknowledgments

The authors acknowledge the Water Research Commission (WRC), the Department of Science and Technology (DST), and the National Research Foundation (NRF) of South Africa for funding. STLH holds the South African Research Chair in Bioprocess Engineering (UID 64778) which has funded this research, as has the WRC/K52392. RJH is funded by a DST/NRF Research Career Advancement fellowship (UID 91465). TM acknowledges studentship support from the NRF (UID 101469). The central analytical facility (CAF) at Stellenbosch University is acknowledged for technical support for biofilm analysis in terms of sulphur content. Ms Miranda Waldron at the UCT imaging and analysis unit undertook the scanning electron microscopy imaging.

